# Long-read sequencing shows complex structural variants in tumor-stage mycosis fungoides

**DOI:** 10.1101/2023.07.03.547529

**Authors:** Carsten Hain, Rudolf Stadler, Jörn Kalinowski

## Abstract

Mycosis fungoides is the most common cutaneous T-cell lymphoma. Recurrent copy-number variations are the main unifying mutations in this disease, but to date, a comprehensive analysis of occurrence and type of structural variants responsible for these copy-number variations remains elusive. In this study, we used Oxford Nanopore Technologies long-read sequencing to elucidate the highly rearranged genomic landscape of five mycosis fungoides samples. We show the occurrence of multiple classes of simple and complex SV and analyze the extend of tumor suppressor gene deletion by complex SVs. Furthermore, leveraging long-read data, we inferred the genomic structure of a chromothripsis event. Our findings highlight the potential of long-read sequencing as a powerful tool for comprehensive genomic analysis in mycosis fungoides.

## Introduction

Mycosis fungoides (MF) is the most common form of cutaneous T-cell lymphoma. In early stages, it presents as fixed inflammatory lesions in the skin. One fourth of patients progress to advanced stage with possible tumor formation and blood involvement (Girardi et al., 2004; Dummer et al., 2021; Krejsgaard et al., 2017; Scarisbrick, 2022). Genetic analysis of MF and the probably related Sézary syndrome shows recurrent copy number variations (CNVs) as driver events, significantly outranking point mutations in number and importance (Choi et al., 2015; Park et al., 2017; Bastidas Torres et al., 2018). Most genetic data was generated with whole-exome sequencing, whose detection power is excellent for point mutations (Miura et al., 2021; Lier et al., 2018), problematic for CNVs (Gabrielaite et al., 2021; Zhao et al., 2020) and almost completely blind to structural variants (SVs) which are causal for CNVs (Cortés-Ciriano et al., 2022; Belkadi et al., 2015). More recent studies on MF genetics used whole-genome sequencing (WGS) and identified SVs as well as commonly rearranged genes, but the main focus remained on the analysis of CNVs (Bastidas Torres et al., 2018; Park et al., 2021).

Comprehensive large-scale analysis of mutations identified SVs as common drivers in cancer and detected different pattern and prevalence of SVs between cancer types. This included simple SVs but also further characterized multiple types of clustered rearrangements forming complex SVs (Li et al., 2020). Complex SVs have far reaching functional consequences, often leading to one or more alterations such as fusion genes, altered gene expression due to acquisition or rearrangement of gene regulatory elements, gene disruption or deletion and altered 3D structure of the genome (The PCAWG Consortium, 2020; Anderson et al., 2018; Cortés-Ciriano et al., 2020; Jeong et al., 2022; Stephens et al., 2011; Wang et al., 2020; Peifer et al., 2015; Cosenza et al., 2022; Yi and Ju, 2018). The combined effect of a complex SV might lead to a jump in the evolutionary landscape greatly enhancing a cancer cell’s fitness and potentially leading to cancer progression (Korbel and Campbell, 2013; Baca et al., 2013; Vendramin et al., 2021).

Further elucidation of cause and effect of complex SVs will need correct reconstruction of their genomic structure. Recent advances in sequencing technology, e.g., regarding accuracy and throughput, made especially Oxford nanopore long-read sequencing well suited for this task. Long-reads show more confident mapping in low complexity and repetitive regions and an improved sensitivity of SV detection compared to short-read sequencing. Furthermore, long-reads ease phasing and allow combined genetic and epigenetic reconstruction of complex genomic regions (Chaisson et al., 2015; Sanchis-Juan et al., 2018; Mahmoud et al., 2019; Chaisson et al., 2019; Coster et al., 2021; Rausch et al., 2023). In addition, long-read RNA sequencing can contribute to inferring the effects of SVs by accurate detection of fusion transcripts across multiple translocations (Tang et al., 2020; Davidson et al., 2022).

## Materials and methods

### Sample selection and DNA purification

Punch and spindle biopsies of MF tumor stage patients were obtained with written consent. DNA was isolated using the DNeasy Blood & Tissue Kit (Qiagen, Hilden, Germany). In addition, sequencing data from the sample published in Hain et al., 2022, including whole-genome PCR library data and adaptive sampling targeted long-read data was added to this project.

### Short-read exclusion

Long DNA fragments were isolated from the samples using Short Read Exclusion XS kit (Pacific Bioscience, Menlo Park, California, USA). The sample MF4 was an exception where the standard Short Read Exclusion kit was used.

### Nanopore whole-genome sequencing

Nanopore sequencing was carried out using the SQK-LSK114 kit (Oxford Nanopore Technologies, Oxford, UK) with a R.10.4.1 flowcell (Oxford Nanopore Technologies, Oxford, UK) on a PromethION 2 Solo (Oxford Nanopore Technologies, Oxford, UK) at 400 bps. Basecalling was done with Guppy 6 (Oxford Nanopore Technologies, Oxford, UK) using the super accurate model with 5mC detection (dna_r10.4.1_e8.2_400bps_modbases_5mc_cg_sup).

### Long-read data analysis

Passing (Q≥10) sequencing data was mapped from unmapped bam file onto T2T CHM13-v2.0 with minimap2 2.24 (Li, 2018) using the -y, -Y, and --MD parameters. For sorting, indexing and bam2fastq conversion samtools 1.13 was used (Danecek et al., 2021). Coverage profiles were made with samtools bedcov using 50 kb bins. Short variants were called with clair3 1.0.1 (Zheng et al., 2022), and after filtering for QUAL≥5 and number of supporting reads > 2, reads were phased with whatshap 1.7 (Martin et al., 2016). Sequencing statistics were gathered using samtools stats, whatshap and NanoPlot (Wouter and Rademakers, 2023).

Structural variants were initially called using delly 1.1.6 (Rausch et al., 2012), svim 1.4.2 (Heller and Vingron, 2019), sniffles2 2.0.7 (Smolka et al., 2022) and cuteSV 1.0.8 (Jiang et al., 2020) with standard or recommended parameters with the following supplements: For sniffles2, the parameters --non-germline and --minsupport 2; for svim, --max_sv_size 200000000; and for cuteSV; --min_support 2 and --max_size 200,000,000. Variant alleles above 500 bp were changed into N for cuteSV. Calls with less than two reads were removed from all vcf files. For filtering, a panel of variant calls from false positive and/or common SVs was be created. Therefore, the filtered vcf files were combined using SURVIVOR 1.0.7 (Jeffares et al., 2017) as a union (SURVIVOR merge 50 1 0 0 0 30), thus combining all calls and only filtering duplicates. The union calls of all 5 samples were intersected using SURVIVOR and calls present in three or more samples were retained (SURVIVOR merge 50 3 0 0 0 30), thus yielding a vcf file containing artifact and/or common SVs. Additionally, the per-caller vcf files of individual samples were intersected and calls present in two or more files were retained (SURVIVOR 50 2 0 0 0 30). This intersection was further filtered by removing calls with less than four supporting reads, calls also present in the above created false positive and/or common SV file and calls present in a lift-over of structural variants from the database of genomic variants (MacDonald et al., 2014) as obtained from UCSC genome browser (Karolchik et al., 2004). The remaining SVs calls were filtered for length above 5 kb and false-positivity due to long-insertion using a custom python script and were finally inspected manually.

### Nanopore RNA sequencing and analysis

RNA was isolated from a skin biopsy using the QuickRNA Miniprep Plus kit (Zymo Research, Irvine, California, USA). Sequencing library was prepared with the SQK-PCS109 kit (Oxford Nanopore Technologies, Oxford, UK) and was sequenced on a R.9.4.1 flowcell (Oxford Nanopore Technologies, Oxford, UK) on a GridION (Oxford Nanopore Technologies, Oxford, UK). Basecalling was done with Guppy 6 (Oxford Nanopore Technologies, Oxford, UK) using the super accurate model. Data was mapped to the reference genome T2T CHM13-v2.0 using minimap2 with the -x splice parameter (Li, 2018) and inspected manually for reads mapping into known exons and spanning SVs.

### Copy number assignment

Number of bases in segments between called SVs was calculated with samtools bedcov. From this, the coverage was calculated, and segments were segmented in copy numbers by manually chosen threshold values. Prior tumor-normal whole-exome sequencing (data not shown) indicated whole-genome duplication for sample MF1 because of the occurrence of multiple even copy number states and the presence of somatic point mutations in low copy number segments. For selected segments, copy number was assigned manually in accordance with coverage and mapping data.

### Gene level analysis

More detailed analysis was carried out for genes from the COSMIC gene census (Sondka et al., 2018) or genes recurrently mutated in CTCL (Park et al., 2017; Bastidas Torres et al., 2018). For genes from this set that were amplified or deleted in at least two samples with one sample showing a focal CNV, the copy number state in all samples was gathered, and if the SV causing the CNV consisted of three or more likely linked SVs or clearly matched a complex SV pattern, the CNV was classified as complex. Short variants were annotated with Ensembl variant effect predictor (McLaren et al., 2016). From all missense variants, variants in selected genes with more than three occurrences in COSMIC and an population allele frequency below 0.1 according to gnomad 2.1.1 (Karczewski et al., 2020) were gathered manually.

### Determination of tumor fraction

After copy number assignment, mean coverage of large (>100 kb) segments of individual copy number states was gathered and a linear fit (coverage = a·CN+b) was computed. Tumor fraction was calculated as b/(a·2+b).

### Methylation

Locations of CpG islands was downloaded from UCSC genome browser (Karolchik et al., 2004) and methylation frequency per region was calculated with modbamtools 0.4.8 (Razaghi et al., 2022). CpG island at tumor suppressor gene promoter sites were inspected manually. For the methylation analysis of SV, the SV supporting reads in the bam file were grouped into a separate phase and supplementary reads were flagged as primary. From these data, the methylation frequency of individual CpG sites per phase was determined using modbam2bed 0.9.4 (Oxford Nanopore Technologies, Oxford, UK) and either plotted using a custom script or manually inspected in IGV (Robinson et al., 2011).

### Homology detection

Homologous sequences at breakpoints were detected using a custom script. Therein, iteratively longer *k*-mers from the reference sequence 10 bp around each breakpoint were computed and both groups were compared for overlaps, with one mismatch allowed.

### Data visualization

Data was visualized using Circos (Krzywinski et al., 2009), IGV (Robinson et al., 2011), Geneious Prime 2022 (Biomatters, Auckland, New Zealand) or with custom scripts.

### Code availability

Custom python scripts used for filtering false positive SV due to insertions and automated assembly of haplotypes containing multiple SVs are available at https://github.com/carstenhain/.

## Results

### ONT sequencing and SV calling of five tumor-stage MF patients

Skin biopsies of four MF patients with tumor stage disease (MF1-MF4) were gathered, DNA was isolated, and samples were sequenced on a ONT PromethION P2. Furthermore, ONT data from a plaque stage patient already published (Hain et al., 2022) was included in this study (MF5). Patient data and general sequencing metrics are listed in Table 1. Tumor cell fraction varied between samples, with only 36 % tumor cells in sample MF4 and 88 % tumor cells in sample MF3.

**Table 1:** Patient data and sequencing metrics for five MF patients including tumor stage, sex, tumor fraction, read coverage, read N50, and phaseblock NG50.

| Patient ID | MF Tumor stage | Sex | Tumor fraction | Coverage | Read N50 [kb] | Phaseblock NG50 [kb] |
| --- | --- | --- | --- | --- | --- | --- |
| MF1 | T3 | F | 0.67 | 34.3× | 9.4 | 333.1 |
| MF2 | T3 | F | 0.83 | 22.7× | 6.0 | 25.7 |
| MF3 | T3 | M | 0.88 | 19.9× | 8.7 | 95.2 |
| MF4 | T3 | F | 0.36 | 18.5× | 13.8 | 249.6 |
| MF5 | T1b | M | 0.63 | 24.5× | 3.2 | 10.2 |

Samples were sequenced to depth between 18.5× and 34.3×. Read length N50 varied between 3.2 and 13.8 kb, while phaseblock NG50 ranged from 10.2 to 333.1 kb. In both metrics, the PCR-based library from Hain et al., 2022 was considerably shorter than the native libraries. Phaseblock length differed between samples and showed the expected combined dependance on coverage and read length.

SV detection was carried out using the consensus from four SV callers (Smolka et al., 2022; Heller and Vingron, 2019; Jiang et al., 2020; Rausch et al., 2012) and subsequent filtering. In detail, filtering included multiple steps: Removal of sequencing artifacts and common germline artifacts by subtracting SVs present in more than three samples or in the database of genomic variants (MacDonald et al., 2014). Further removal of germline SVs was achieved by filtering SVs smaller than 5 kb. This is a tradeoff in which a disproportionate number of the mostly small germline SVs are filtered out, but small somatic mutations are also lost in the process. Despite the focus on removing germline SVs, rarer larger events, for example due to the deletion or insertion of the 6 kb L1 retrotransposon (Cordaux and Batzer, 2009), might remain. Final filtering steps involved removal of false positive SVs due to mis-mapping of reads at insertion sites using a custom script and the manual checkup of the resulting SVs.

Copy number and SV analysis in all five samples (Figure 1a-e) revealed a shattered copy number landscape and various large (>5 kb) SVs. The number of SVs per sample ranged from 129 to 281 (Figure 1f), which is much higher than the previously described mean of 39 SVs in tumor-stage MF patients (Bastidas Torres et al., 2018). As four of five samples showed tumor fraction above 63 %, mean variant coverage should be sufficiently above the chosen cutoff of four. This contrasts sample MF4, which has a considerably lower tumor fraction. Therefore, the mean variant coverage should be around 3.3 (mean coverage · tumor fraction · 0.5), which is lower than the chosen cutoff. Therefore, a sizeable fraction of SVs was probably not detected in sample MF4. Furthermore, sample MF5 is a PCR-based library resulting in dropout of individual regions and an inflated coverage due to PCR duplicates, making comprehensive calling of variants challenging even in this high tumor fraction sample.

**Figure 1:**
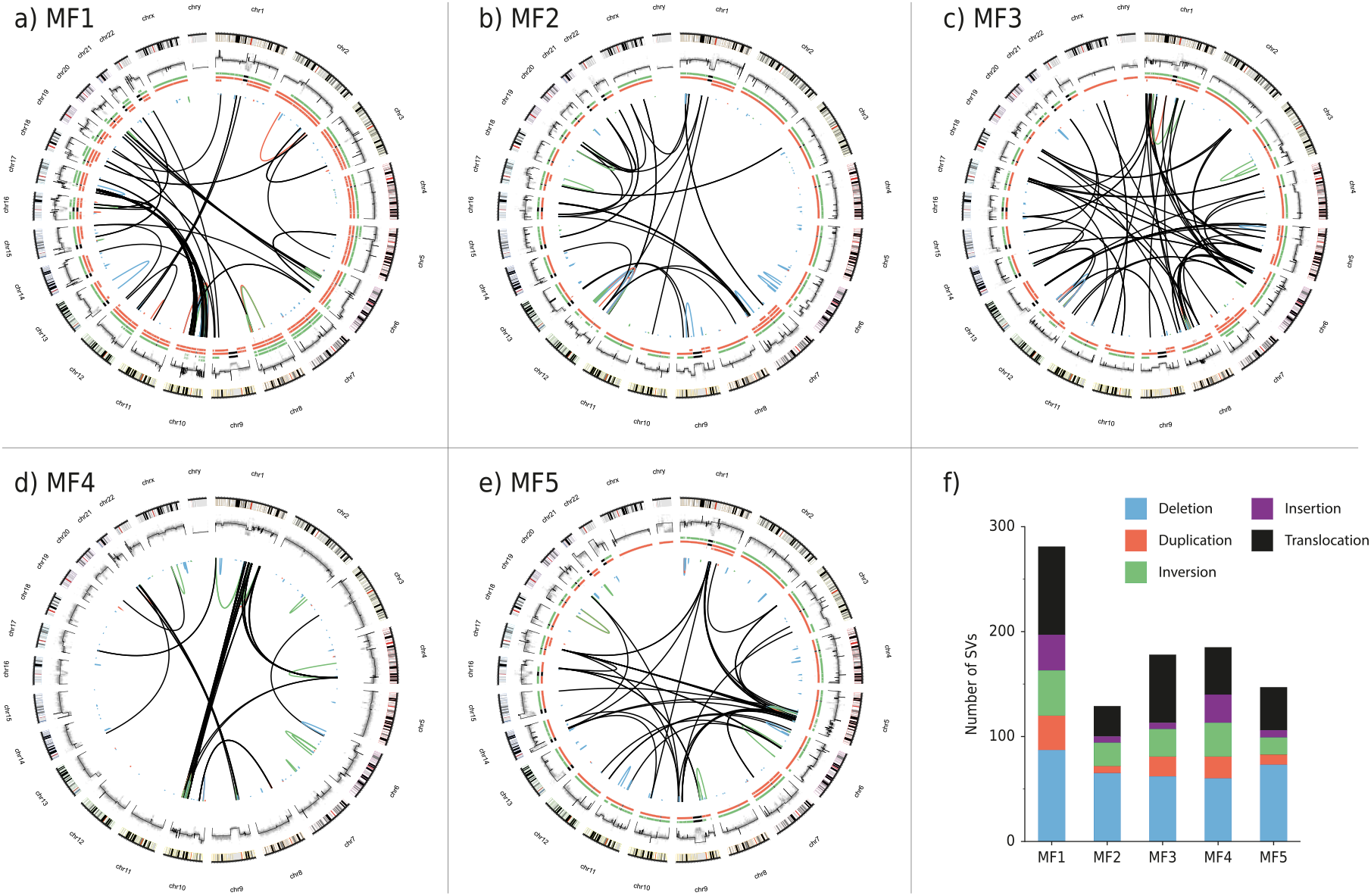
Genomic landscape of five MF patients. **a-e)** Circos genome plots showing the following, described from the outside to the inside: the chromosomes, raw and segmented coverage, assumed tumor cell copy number for the major (red) and minor (green) allele and structural variants (>5 kb) colored by type as indicated in f). Black segments denounce regions omitted from analysis. Assignment of copy number was not attempted from sample MF4 due to low tumor fraction. **f)** Number of structural variants (>5 kb) found in each sample.

Detection of homology in a 20 bp window around breakpoints failed most of the time and was found only for shorter (<20 kb) deletions if any (Figure S1). This indicates a dominant homology-free repair mechanism, probably based on microhomology or the non-homologous or mismatch-mediated end joining pathway (Li et al., 2020).

With the exception of sample MF3, deletions are the most frequently found SV type, followed by translocations. In MF3, this finding is reversed. Insertions are the rarest SV type and the insertions found are often about 6 kb long, probably indicating L1 retrotransposon insertions. The detection of insertions longer than the average read length is increasingly challenging, as reads completely spanning the insertion become sparse (Mahmoud et al., 2019). Therefore, the low number and size of insertions must be seen with care.

All samples show extensive copy number alterations, with MF1 showing signs of a whole-genome duplication and, for instance, four samples (MF1, 2, 3, 5) showing chr1q and chr7 amplification.

The SVs are not evenly distributed on the genome and some chromosomes lack any large SV or translocation while each sample has at least one or more, often connected dense clusters of SVs. Defined examples are the chr10-chr17 cluster of MF1 with 98 SVs and the chr1-chr10 cluster of MF4 with 32 SVs. These clusters often show oscillating copy number patterns with chr10 in MF1 oscillating between copy number of 2 and 4, while a 18 Mb region with 23 SVs on chr8 in MF3 oscillates between copy numbers 2 and 3.

Another frequent pattern of SVs are chains of balanced translocations. Here, multiple occurrences of inwards facing pairs of SVs are connected to each other. SVs of individual pairs are often in close proximity but may be separated by a few kb to multiple Mb, leading to deletions between adjacent breakpoints.

### Nanopore sequencing enables simplification and putative reconstruction of a chromothripsis event

The sample MF1 shows a dense occurrence of SVs connecting chr6, chr10 and chr17 (Figure 1a, Figure 2a). In total 105 SVs on chr10 and chr17 as well as translocations between both chromosomes are connecting 104 segments of copy number 4. Furthermore, two translocations between chr6 and chr10 are detected. This cluster shows characteristics of a chromothripsis event, namely the oscillating copy number, the occurrence of breakpoint clusters (Figure S2), the occurrence of all orientations of SVs, the retained heterozygosity on the high copy number segments and the loss of heterozygosity on the low copy number segments (Korbel and Campbell, 2013; Cortés-Ciriano et al., 2020).

**Figure 2:**
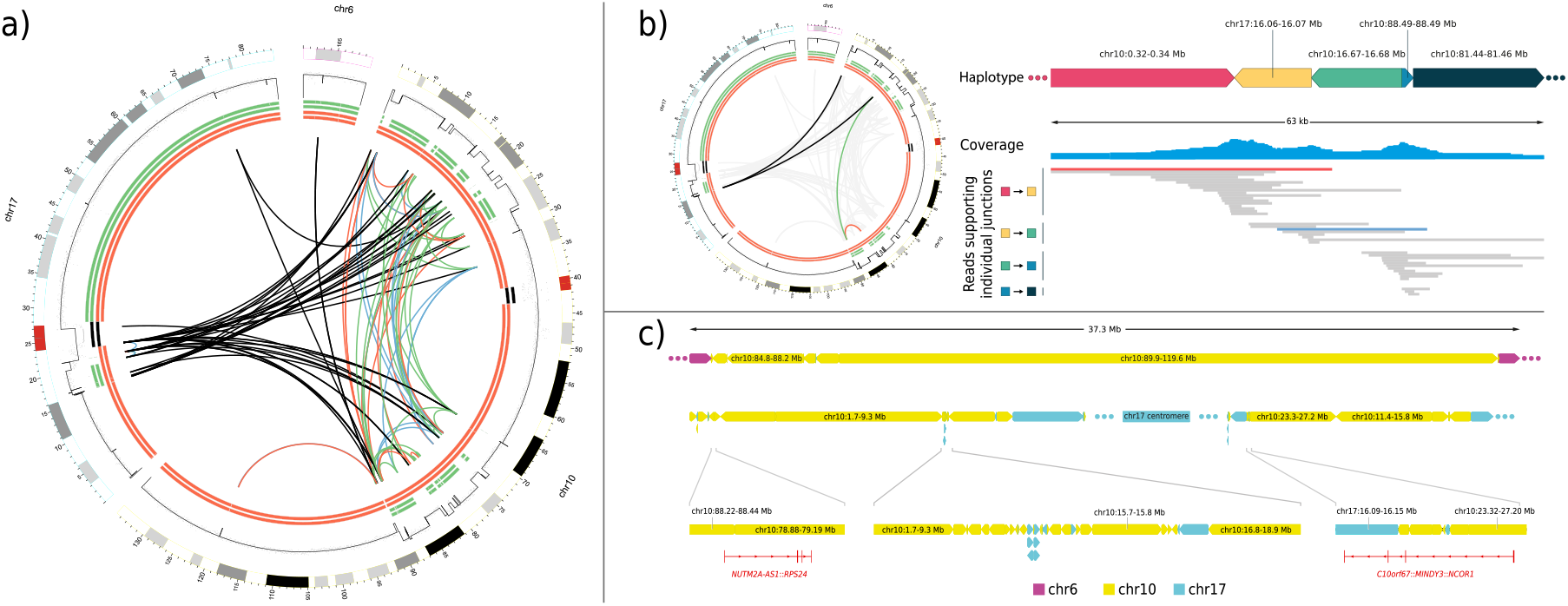
Long-read sequencing simplifies and enables putative reconstruction of a chromothripsis event. **a)** Sample MF1 shows 107 translocations mostly between chr10 and chr17 but also chr6 and an oscillating copy number, which indicates a chromothripsis event. Circos plot is structured as in Figure 1. **b)** Iterative assembly of nanopore reads connects multiple SVs directly. The four translocations highlighted in the Circos plot (left) are connected by long reads and form the depicted haplotype. The iterative process of read selection, assembly and SV calling is depicted with groups of reads initially selected as supporting of one single translocation (indicated by two colored boxes on the left). As the resulting assembly elongates, new translocations are detected leading to selection of a new batch of reads. Exemplary reads connecting more than one SV are colored red and blue. **c)** Putative reconstruction of the chromothripsis event led to three blocks. The first block indicates an insertion of 13 segments from chr10 and 1 segment from chr17 into chr6 (top). The second block starts from the chr10 5’-end, transverses 43 segments from chr10 and 23 segments from chr17 and ends into the chr17 centromere with + direction (middle-left). The third block connects 27 segments from chr10 and 11 from chr17 and faces towards the chr17 centromere on one side and to the chr17 3’-end on the other side (middle-right). A potential connection via the chr17 centromere is indicated. Interesting junctions, either creating fusion genes as shown by long-read RNA-sequencing (bottom-left/right) or a 300 kb stretch concatenating multiple small segments completely spanned by long reads or phasing, are shown (bottom-middle).

The segments of high copy number are partly in range to be spanned completely by individual sequencing reads, enabling the connection of multiple SVs into haplotypes. We therefore devised a script, that starting from a single SV iteratively builds haplotypes connecting multiple SVs. Each iteration works by a) gathering reads supporting the initial SV in the first round and all SVs on this haplotype in later rounds, b) read assembly using lamassemble (Frith et al., 2021), c) alignment of the assembly to the reference genome and detection of additional SVs spanned by the haplotype assembly (Figure 2b). This resulted in 34 subassemblies with sizes between 5.2-71.1 kb (mean 27.8 kb) and 0-22 spanned segments (mean 2.15). Subassemblies connected by completely phased segments were manually connected resulting in 22 subassemblies with sizes between 5.2-606.8 kb (mean 89.6 kb) and 0-33 spanned segments (mean 3.86). In total, this approach simplified the chromothripsis event from 98 SVs to 22 subassemblies only connecting large segments of the same copy number. Further, manual connection of the large segments, neither spanned by individual reads or phasing, by their flanking subassemblies yielded three putative reconstructions (Figure 2c). The first reconstruction includes both translocations to chr6 and indicates that 13 segments from chr10 and 1 segment from chr17 are inserted into chr6 as a 35.3 Mb insertion. The second reconstruction starts from the 5’-end of chr10 and connects 43 segments from chr10 and 23 segments from chr17. It ends inside the chr17 centromere, facing towards the chr17 3’-end. The third segment connects 27 segments from chr10 and 11 segments from chr17. It starts and ends on chr17 and faces towards the chr17 centromere and towards the chr17 3’-end. Therefore, a connection of the second and third segment via the chr17 centromere is possible and would create a chr10-chr17 fusion from 5’-chr10 to 3’-chr17 with the chr17 centromere.

This chromothripsis event leads to widespread deletion of one allele and loss of heterozygosity on chr10 and chr17, including numerous tumor suppressor genes (TSGs) like *TP53*, *ZEB1*, *NFKB2*. In conjunction with a deletion on the non-rearranged allele, this event leads to a homozygous *FAS* deletion. Furthermore, this event forms fusion transcripts at its junctions between different segments. The *NCOR1*::*MINDY3*::*C10orf67* and *NUTM2A*-*AS1*::*RPS24* fusions were detected by long-read RNA sequencing. Prediction of the functional effects of these fusion genes is challenging, but individual components, like the ribosomal protein 24 (RPS24) can act as oncoproteins, e.g. in different colon and prostate cancer cell lines (Kazerounian et al., 2016; Li et al., 2023; Wang et al., 2015).

### Different complexities of connected SVs lead to tumor suppressor gene deletion

In each sample, there are several concatenated occurrences of the pattern "breakpoint-deletion-breakpoint", whereby the length of the deletion varies greatly between 0 bp and multiple Mb. These chains can link two to above 10 of the patterns on one or more chromosomes (Figure 3, Figure S3). Based on size, number of affected chromosomes and size of the deletions between breakpoints, these chained patterns either resemble the chromoplexy class of complex SVs or the class of local n-jumps (Li et al., 2020; Baca et al., 2013; Yi and Ju, 2018).

**Figure 3:**
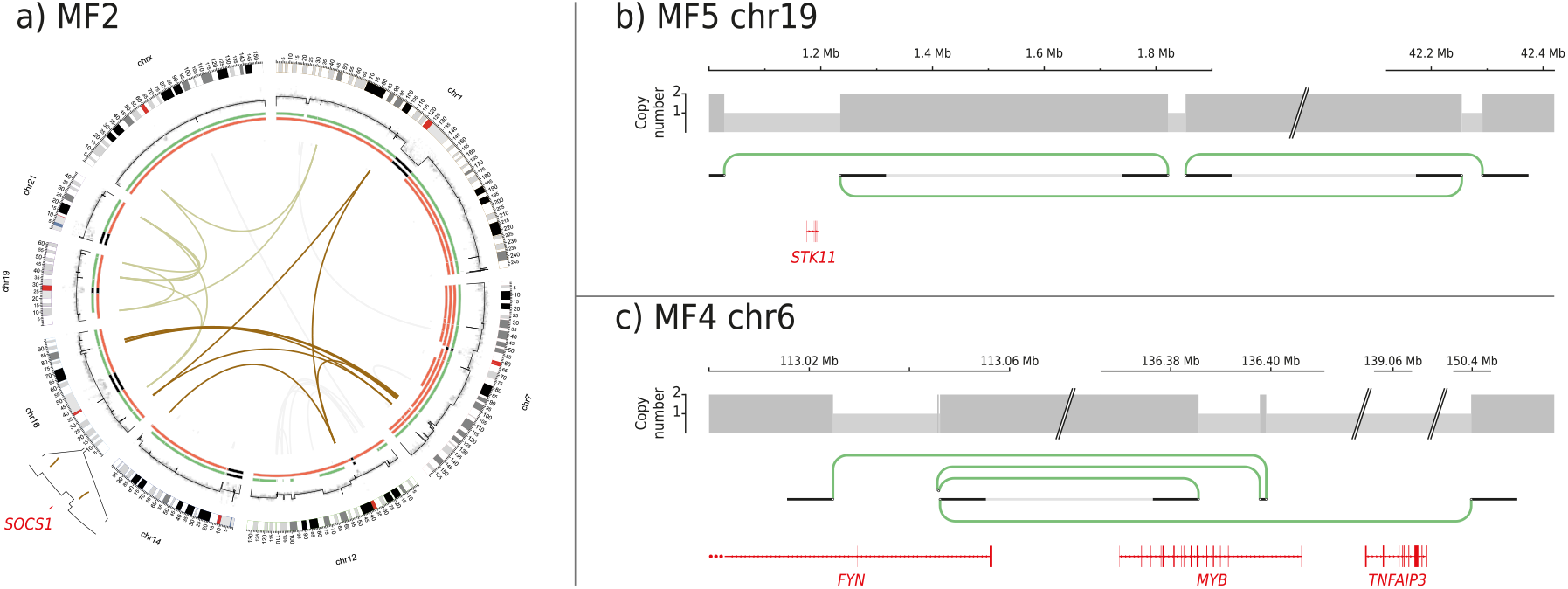
Interconnected chains of SVs of different complexity led to TSG deletion and putative formation of fusion genes. **a)** Two instances of chromoplexy (olive-green and brown) are shown for sample MF2. The olive-green chromoplexy is a completely closed chain of 11 SVs on 5 chromosomes, while the brown chromoplexy connects SVs on 5 chromosomes and ends at the terminal truncation of chr14 and inside the cluster on chr12. The brown chromoplexy deletes one allele of the JAK-STAT suppressor *SOCS1*. SVs in the Circos plot are greyed out if they are not part of one of the depicted chromoplexy events. The gene location (red), coverage profile and exact location of SVs at the *SOCS1* locus is shown in detail. Circos plot is structured as in Figure 1. **b)** Local 3-jump on chr19 in sample MF5 leading to *STK11* deletion and **c)** local 4-jump on chr6 in sample MF4 leading to *TNFAIP3* deletion and the putative formation of a *MYB*::*FYN* fusion. Both plots show copy number (grey boxes) and SVs colored in green to indicate them as inversion type. The position of the areas shown is labelled at the top. Compression of the genomic area is indicated by skewed lines. Exon structure of genes of interest are shown in red.

Sample MF2 shows two distinct chromoplexy events (Figure 3a), with one completely closed chain on chr1, chr16, chr19, chr21 and chrX. The other event is chaining between chr1, chr7, chr12, chr14 and chr16 and ends next to a terminal deletion of chr14 on one end and leads into the cluster of SVs on chr12 on the other end. This chain deletes one allele of the TSG *SOCS1* on chr16 in one of its focal deletions between translocations. The other allele of *SOCS1* in this sample is deleted by a simple deletion. The *SOCS1* deletion is a well-known TSG deletion in MF and might abolish negative feedback of JAK-STAT signaling. It was found in tumor and corresponding early stages of MF patients, and the combination of simple deletions and deletions due to translocations for biallelic *SOCS1* deletion was also demonstrated (Bastidas Torres et al., 2018). Additional notable deleted genes due to this chain are the oncogenes *SMO* and *SND1* on chr7. If these genes are mutated, they are generally affected by activating mutations, but in cases of acute myeloid leukemia and esophageal carcinoma, *SND1* is rare but recurrently deleted (Cui et al., 2020).

Further chromoplexy events were observed e.g., in sample MF3 leading to deletion of *PRDM1*, *RB1* and *DLEU2*. In addition, the dense clusters connecting chr1 and chr10 in sample MF4 as well as the dense cluster on chr5 in MF5, connected to multiple other chromosomes, show multiple occurrences of individual chains of SVs, thus resembling chromoplexy. However, comprehensive variant calling in these samples is difficult due to low tumor fraction or library type leading to incomplete SV calling and hindering complete resolution of genomic structure in these areas.

Examples of local n-jumps are the local 3-jump on chr19 for sample MF5 leading to *STK11* deletion (Figure 3b) and the local 4-jump on chr6 for sample MF4 leading to *TNFAIP3* loss as well as the putative formation of a *MYB*::*FYN* fusion (Figure 3c). The deletion of *STK11* and *TNFAIP3* were recurrently found in MF (Bastidas Torres et al., 2018; Park et al., 2017). *MYB* fusions are recurrent in adenoid cystic carcinoma, where the *MYB*::*NFIB* fusion causes MYB overactivation (Wagner et al., 2022).

### Variant aggregation shows recurrent deletion of TSGs by complex SVs

As discussed, the MF tumor samples are highly rearranged and contain multiple complex SVs. Aggregation of the copy number state of genes from the COSMIC gene census and genes recurrently affected by CNVs in other CTCL studies (Park et al., 2017; Bastidas Torres et al., 2018) reveals the extent of the influence of complex SVs on the copy numbers landscape of MF (Figure 4). Here, only genes affected by CNVs in two samples with at least one focal CNV are shown, and the allelic state is resolved. Depending on the type of SV leading to loss of an allele, deletions are divided into simple deletions and complex deletions. A deletion is termed complex if it results from two or more connected SVs (Liu et al., 2011). One example is the mentioned homozygous deletion of *SOCS1* in MF2 due to one simple deletion and one chromoplexy event. In total, 35 alleles of known TSGs are deleted by simple deletions and 17 alleles are deleted by complex events; thus, complex SVs account for one third of all deletions in this set of genes. This again illustrates the importance of complex SVs in the mutational landscape of MF.

**Figure 4:**
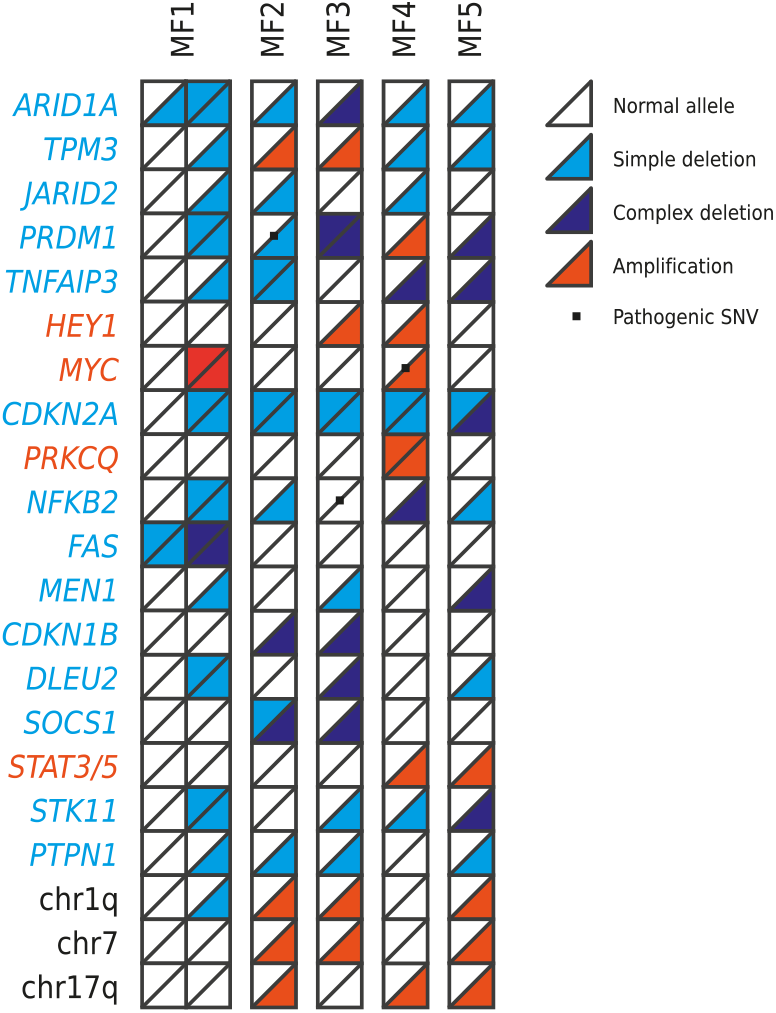
Genes recurrently affected by CNVs in five MF samples. The copy number state of various TSGs (blue), oncogenes (red) or broad chromosomal regions (black) is indicated for each sample. Single alleles are indicated as triangles. Normal state is colored white and amplified alleles are colored red. Alleles deleted by simple SVs are colored light blue, and alleles deleted due to complex SVs are colored dark blue. Pathogenic SNVs are shown as black dots. A copy number of four is assumed as the ground state of the whole-genome duplicated sample MF1.

Next to SVs and CNVs, various SNVs and different methylated promoters were detected using this dataset. As no patient matched normal data was available, a complete differentiation between germline and somatic SNVs is difficult. But several SNVs at known hotspots in cancer genes such as *JAK3*, *NFKB2*, *MYC* and *CDKN1B* were found (Figure 4). Furthermore, the samples show a varying degree of different methylated promoters of cancer genes, including hypermethylated promoters of TSGs like *CDH1*, *PTPRB*, *PTPRT*, *SFRP4* (Figure S4).

In addition to highlighting the importance of complex SV in the mutational landscape of MF, this study indicates *PRDM1* as a recurrently lost TSG in MF. This gene is homozygously inactivated in sample MF2 by a point mutation and a simple deletion and in MF3 by two complex SVs. Additionally, MF1 and MF5 show heterozygous deletion of *PRDM1*. PRDM1 is a zinc-finger containing transcriptional repressor that regulates genes involved in cell proliferation and survival and is involved in B- and T-cell development (John and Garrett-Sinha, 2009; Boi et al., 2015). Recurrent biallelic inactivation of PRDM1 were found in T-, B- and NK-cell lymphoma (Boi et al., 2013; Boi et al., 2015; Küçük et al., 2011; Mandelbaum et al., 2010; Xia et al., 2017) and CD30+ lymphoproliferative disorders (Argyropoulos et al., 2020). Until now, *PRDM1* loss was detected only in one advanced stage of MF (Argyropoulos et al., 2020) and was not indicated in a larger MF/CTCL data collection (Park et al., 2017; Park et al., 2021; Iyer et al., 2020).

## Discussion

Unifying genetic knowledge of MF is lacking, with multiple studies identifying various SNVs, but nonetheless, only limited consensus and recurrence between samples exist. More comprehensive overlaps were detected on the occurrence of large alterations affecting chromosomal structure and the genomic copy number of individual genes or large segments. But due to prevalent use of array (Gug et al., 2019) and whole-exome sequencing data (Park et al., 2017), this information is mostly present only as CNV information, meaning amplification or deletion of genes and chromosomal regions without detecting the underlying SVs. Use of WGS already indicated complex SVs in CTCL cases (Choi et al., 2015) and increasing use of WGS in studies showed their occurrence in MF (Bastidas Torres et al., 2018) as well as detected genes recurrently disrupted by translocations (Park et al., 2021). However, a focus on complex SVs in MF is missing so far (Bastidas Torres et al., 2018; Park et al., 2021). To add this information, this study uses long-read sequencing to detect multiple types of complex SVs and shows their impact on known cancer genes, including their frequent involvement on the deletion of TSGs. Each of the five sequenced samples contained at least one complex SV and complex SVs were the cause for one third of deleted TSG alleles.

One sample showed chromothripsis with oscillating copy numbers and 100+ translocations on and between chr10 and chr17 as well as two translocations to chr6. Long-read sequencing simplifies this complex rearrangement by connecting multiple SVs either directly by individual reads or by phase information, ultimately allowing complete putative reconstruction of the shattered segments into one 35.3 Mb insertion into chr6 and a chr10-chr17 fusion chromosome.

Chromothripsis is a pan-cancer phenomenon present in 29 % of cancer samples with large type-specific differences and various characteristics regarding copy number state, number of affected chromosomes and number of SVs. Chromothripsis events connecting multiple chromosomes, as in this sample MF1, could point to simultaneous fragmentation of several chromosomes and/or a previous translocation between chromosomes (Cortés-Ciriano et al., 2020). Chromothripsis was already detected in two SS (Choi et al., 2015) and three MF tumors (Bastidas Torres et al., 2018).

As shown here and elsewhere (Rausch et al., 2023; Cretu Stancu et al., 2017; Miller et al., 2021; Scharf et al., 2022), long-read sequencing enables the resolution of complex genomic rearrangements, thus making them accessible for further analysis, such as mapping of RNA sequencing data or systematic analysis of breakpoint location and potential rearrangement pattern. For the reconstruction in this paper, subassemblies were created by iterative assembly starting from a single SV using a custom script and subassemblies were used to manually connect large segments. Other approaches for the resolution of complex rearranged regions, such as completely manual arrangement of SVs (Miller et al., 2021), targeted phased assembly (Rausch et al., 2023) or the inference of segment orientation and order from mapping data (Mitsuhashi et al., 2020) exist. All these approaches benefit from increasingly long reads to connect more distant SVs and create larger phase blocks. Furthermore, the tumor fraction influences the success of individual methods and e.g., phased assemblies often represent a mixture of the tumor and normal haplotype in low tumor fraction samples. Complementing the ability to reconstruct the genomic loci of complex SV, long read sequencing enables direct detection of fusion transcripts formed by one or even multiple SVs, thus easily unraveling one of the potential effects of complex SVs (Davidson et al., 2022).

Each of the samples showed multiple occurrences of connected pairs of inwards facing SVs with deletion bridges of various sizes, leading to the deletion of TSGs. Some of the larger chains connecting SV pairs on multiple chromosomes resembles chromoplexy, while smaller chains on single chromosomes interspersed with larger deletion bridges resemble local n-jumps. Because both classes can show similar patterns, sequencing based differentiation between them remains challenging and is mainly based on the number of affected chromosomes: Chromoplexy mostly connects multiple chromosomes, while local n-jumps are confined to a single chromosome. Use of the length of deletions as a distinguishing feature is difficult. For example, chromoplexy is mostly copy number neutral with short deletion bridges, but in rare cases, Mb-long deletion bridges were found in chromoplexy chains (Li et al., 2020; Baca et al., 2013; Anderson et al., 2018). Despite similar patterns, both classes have very different formation mechanisms: Chromoplexy arises from a single catastrophic hit which creates multiple DNA double strand breaks, probably at actively transcribed regions. Afterwards, random shuffling of the DNA ends and subsequent ligation, potentially with major loss of adjacent DNA, leads to the observed closed chains of balanced translocations (Baca et al., 2013). In contrast, local n-jumps are explained by local polymerase template switches (Li et al., 2020) which are likely to occur in the context of mismatch-mediated break-induced replication of DNA double strand breaks that arise from replicative stress (Cosenza et al., 2022).

The frequent occurrence and the shown functional impact indicate the importance of complex SV in MF, at least in the tumor stage, and adds complex SVs as players in generating the recurrent CNVs and large-scale genomic aberrations in CTCL. This calls for future genomic analyses of MF to include the characterization of complex SVs regarding type, occurrence, and effect. For this endeavor, this study can only be the starting point, showing the importance and prevalence of complex SVs and the methodical approach to elucidate them using long-read sequencing. Low sample size and the possible overrepresentation of highly rearranged samples makes further statements from this study difficult. Therefore, future studies should direct special interest to use state-of-the-art methodology and include MF samples of various stages, including untreated early-stage MF, to detect stage-specific prevalence of small, large, and complex somatic mutations. Further integration of longitudinal data might help to establish connections between somatic mutations and disease progression, similar to T-cell receptor beta rearrangement quantification (Masson et al., 2018). In this regard, complex SV may play a pivotal role, as they are known to be important drivers of progression in other cancer types (Vendramin et al., 2021; Korbel and Campbell, 2013).

Going forward, integration of classical SNV and CNV data with the analysis of simple and complex SVs, in multiple stages and timepoints, might resolve the still enigmatic genomic causes of MF.

