## Supplementary Figures for "Long-read sequencing shows complex structural variants in tumor-stage mycosis fungoides"

### Supplement

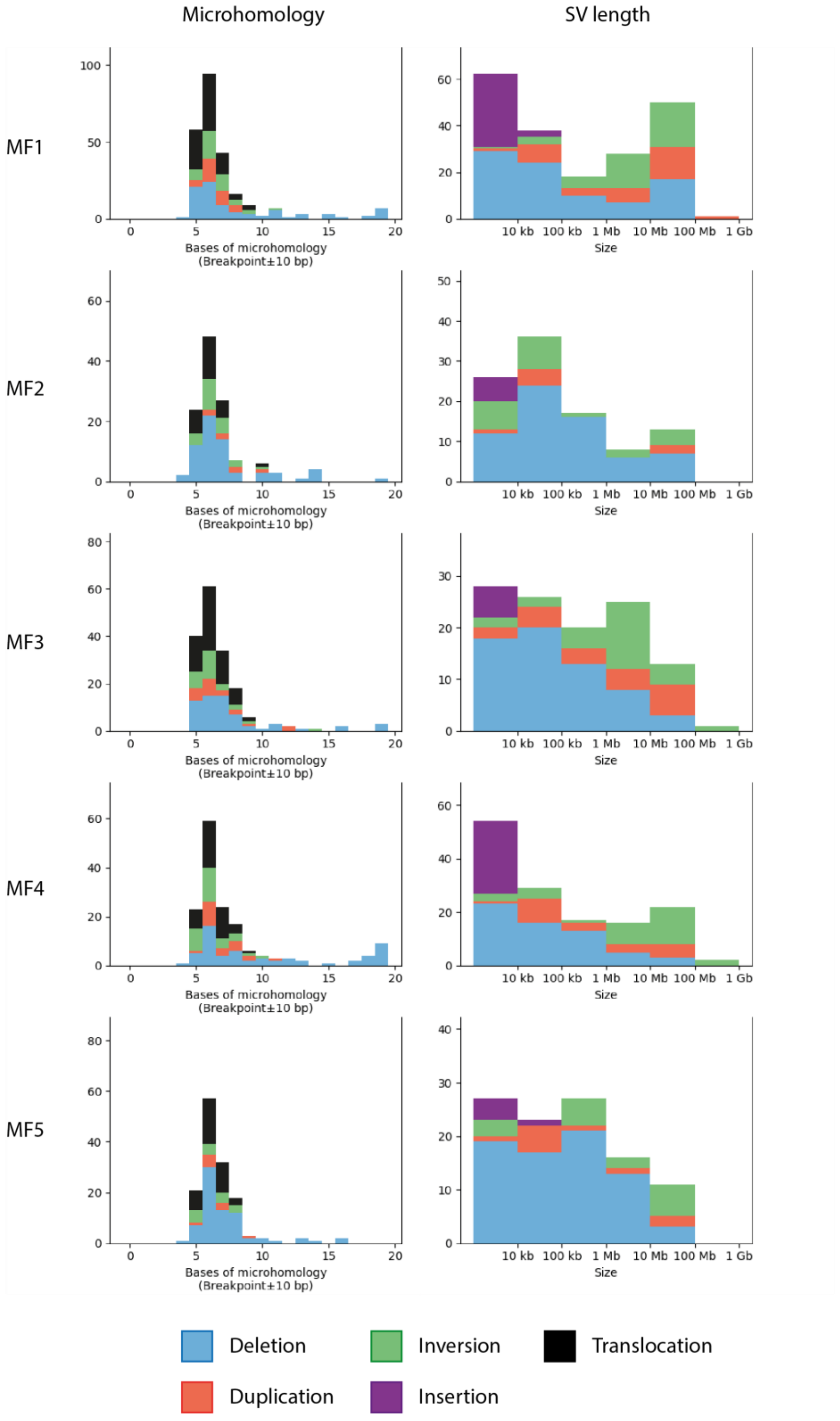


Figure S1: **Homology at breakpoint sites and length of SVs.** a) Size of the largest stretch of homologous bases in an ±10 bp window around both breakpoints. b) Size distributions of SVs per sample. SVs are colored by type.

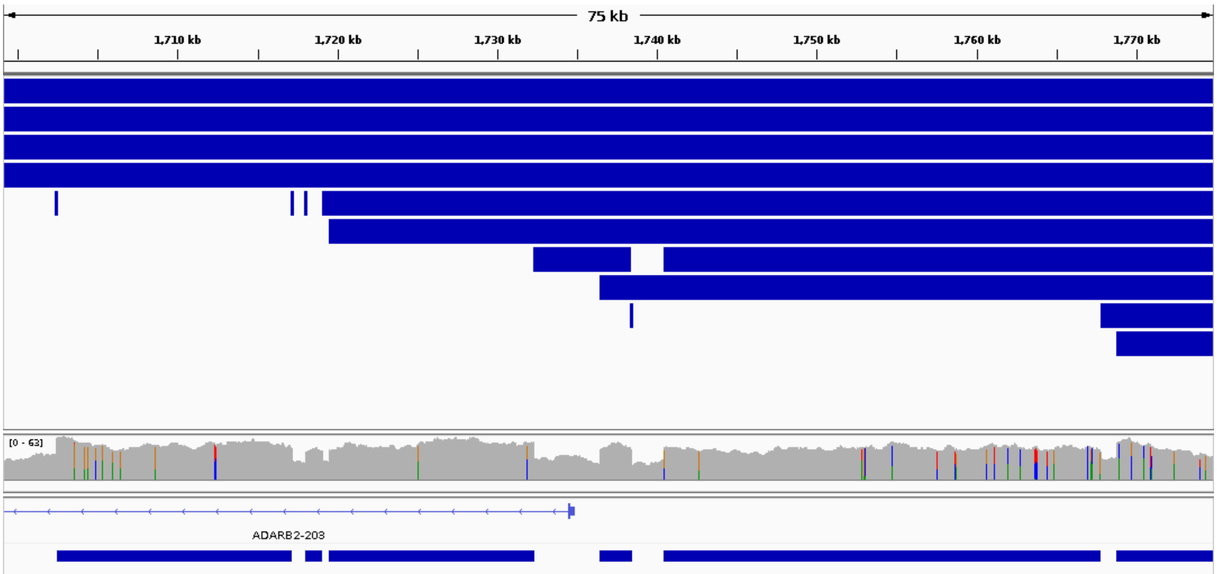


Figure S2: Breakpoint cluster inside the chromothripsis region on chr10 on sample MF1. This 75 kb region contains 11 breakpoints and 10 SVs. From top to bottom: chromosomal position on chr10, SVs (blue boxes), sequencing coverage (grey), and regions with elevated coverage (blue boxes, bottom).

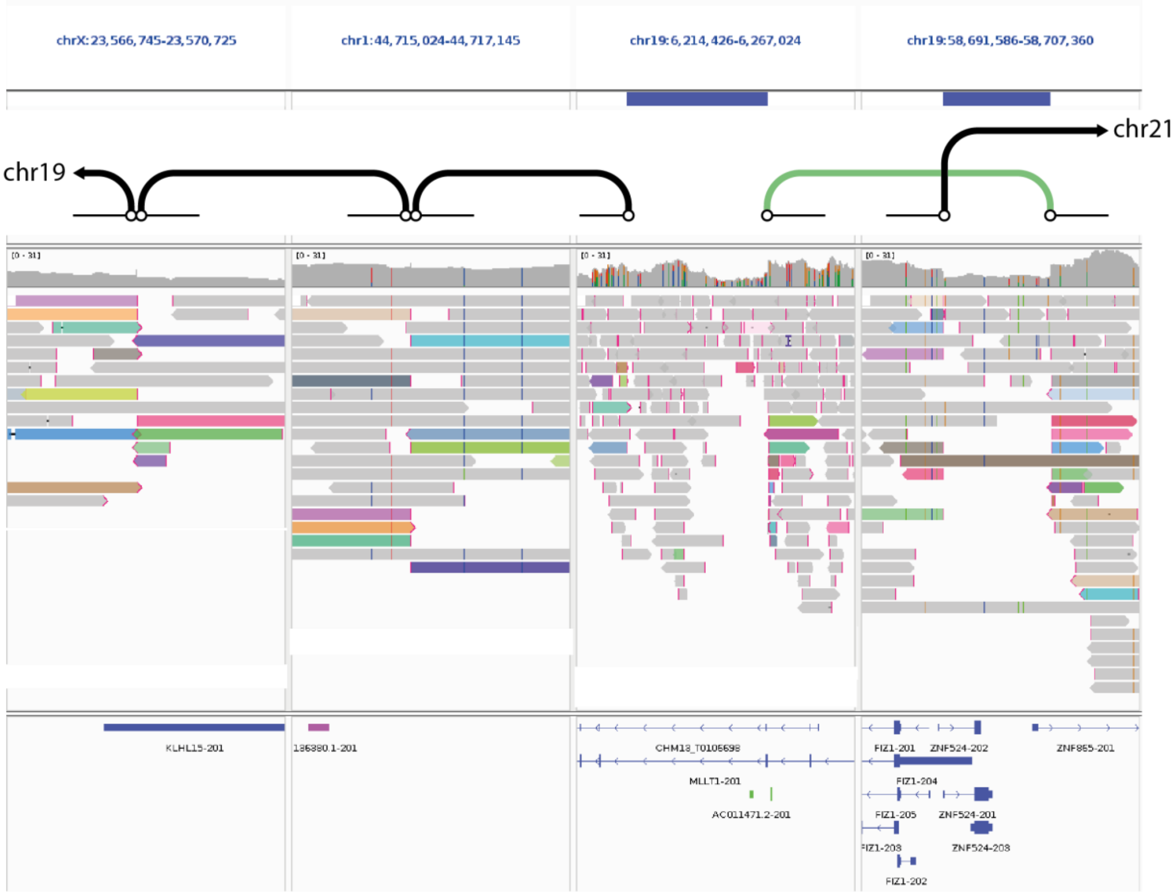


Figure S3: **Excerpt from a chromoplexy chain on MF2 showing multiple connected balanced translocations/SVs on chrX, chr1 and chr19, partly with deletion bridges between both breakpoints (blue regions, top).** Reads with supplementary alignment are colored. SVs are shown as thick lines, with black indicate translocation and green inversion type SVs, connecting two breakpoints (circles) and their adjacent chromosomal segments (thin black lines).

a)

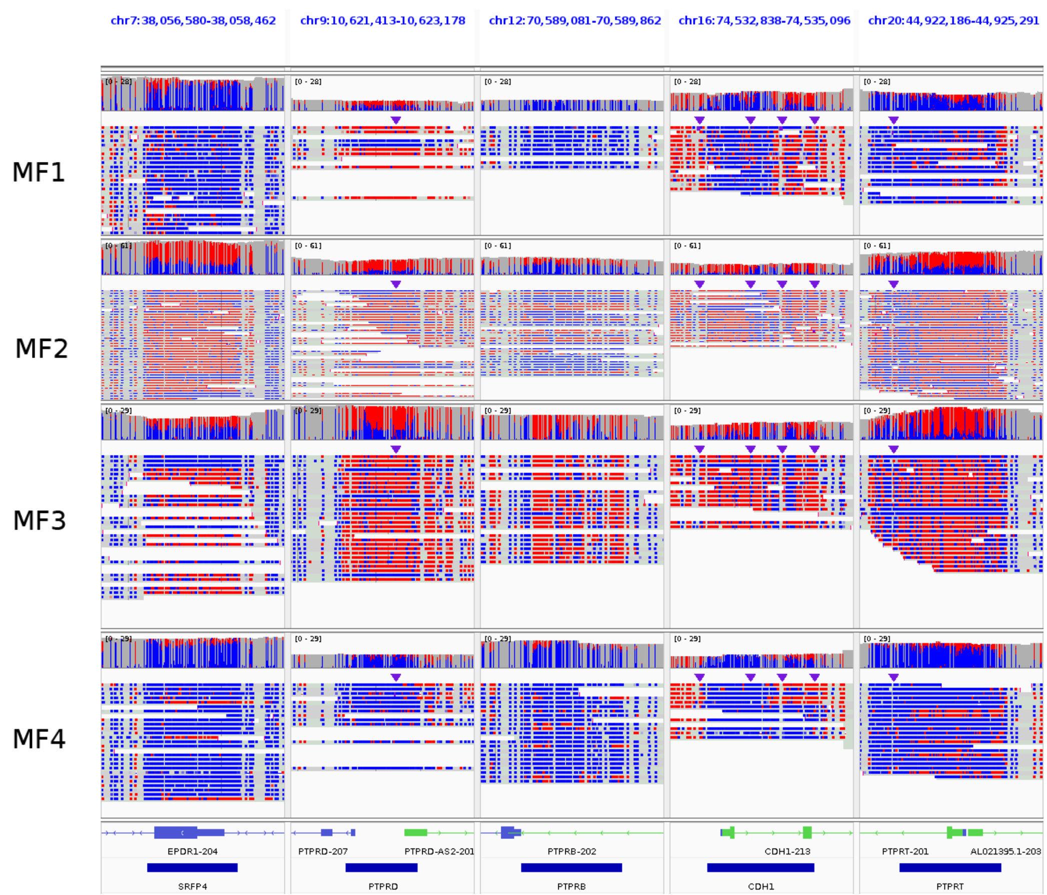


b)

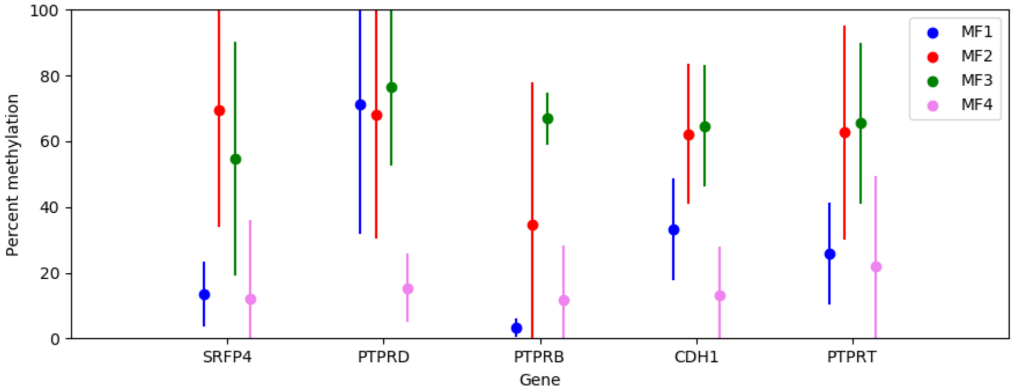


Figure S4: **Different promoter methylation of five TSGs for samples MF1-MF4.** a) IGV view on the CpG island at the gene starts of *SRFP4*, *PTPRD*, *PTPRB*, *CDH1* and *PTPRT* in the samples MF1-MF4. The reads and the coverage are colored according to the methylation state: methylated cytosines are colored red, unmethylated cytosines are colored blue. Other bases, including cytosines outside of the CpG context are grey. b) Percentage of methylated cytosine in the CpG islands from a) as calculated with modbamtools (Razaghi et al., 2022).

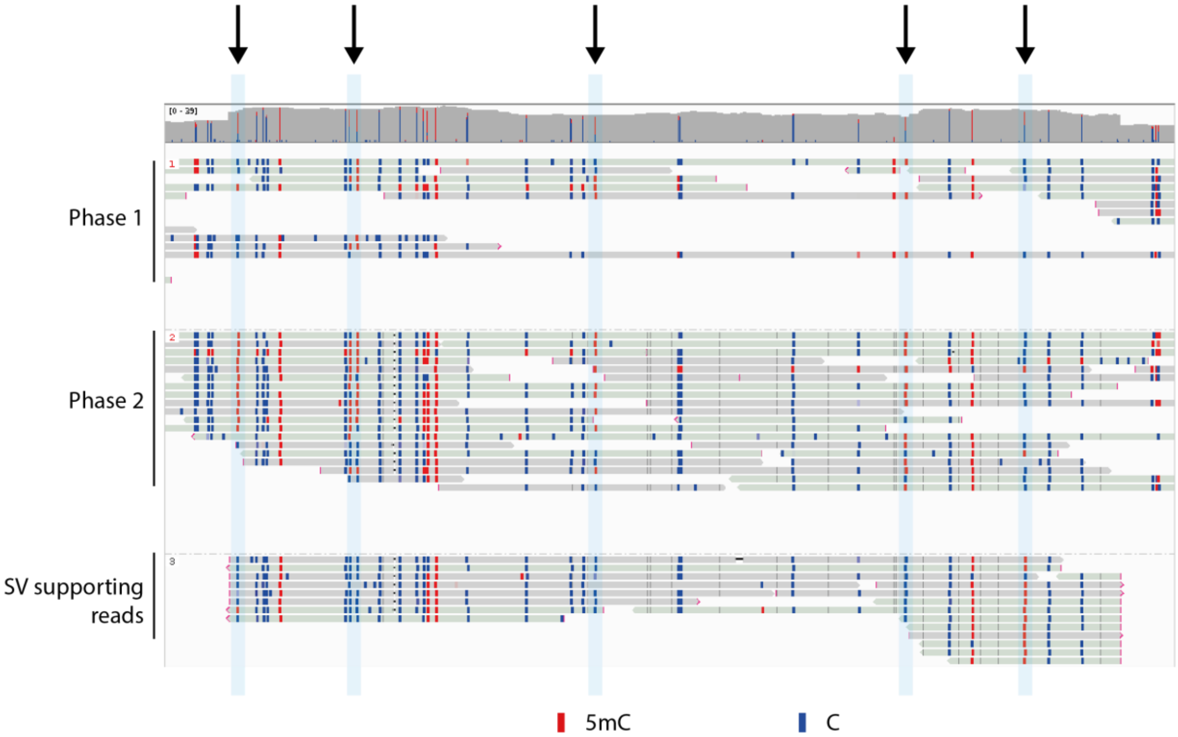


Figure S5: **Different CpG methylation in a complex SV from sample MF3.** Sample MF3 shows multiple connected segments amplified to a copy number of three on chr8. Separation into phases and further grouping of the SV supporting reads show different methylation pattern of the amplified segment. The amplified segment shows SNP pattern similar to phase 2 but shows different methylation at 6 CpG sites (marked in light blue/with black arrows). Inside the CpG context, methylated cytosines are colored red, while unmethylated cytosines are colored blue.
